# Rapid and robust sex determination from ancient enamel proteomes using *protSexInferer*

**DOI:** 10.64898/2026.03.23.713598

**Authors:** Fan Bai, Zhongyou Wu, Song Xing, Qiaomei Fu

**Affiliations:** State Key Laboratory of Animal Biodiversity Conservation and Integrated Pest Management, Institute of Zoology, Chinese Academy of Sciences, Chaoyang District, Beijing, China; Key Laboratory of Vertebrate Evolution and Human Origins of Chinese Academy of Sciences, Institute of Vertebrate Paleontology and Paleoanthropology, CAS, Beijing 100044, China; University of Chinese Academy of Sciences, Beijing 100049, China

**Author notes:** **Correspondence:** (Q. M. Fu). **Author list footnotes** These authors contributed equally (Fan Bai, Zhongyou Wu, Qiaomei Fu).

**Keywords:** paleo-proteomics, sex determination, automated pipeline, AMELY ratio (R_AMELY_), dental enamel

## Abstract

Accurate biological sex determination of ancient remains is critical for archaeological, anthropological, and forensic studies, but remains challenging for morphologically ambiguous and highly degraded endogenous DNA samples. Paleo-proteomics sex identification approaches, targeting sexually dimorphic amelogenin isoforms (AMELX and AMELY), present a promising solution. However, current workflows rely on manual verification of a few specific peptide markers, a process that lacks standardization and is susceptible to false-positive AMELY signals. To overcome these limitations, we developed *protSexInferer*, a lightweight, open-source bioinformatic pipeline for automated sex estimation from paleo-proteomic data. Our method uses the ratio of AMELY-specific peptides to all detected AMELY- and AMELX-specific peptides (i.e., the R_AMELY_ value) rather than the mere presence or absence of AMELY signals for sex classification. We demonstrated that the R_AMELY_ value clearly distinguishes male and female individuals in both reference and independent validation datasets, enabling reliable sex assignments even in cases where conventional intensity-based comparisons (e.g., AMELY-59M vs. AMELX-60) are ambiguous. This ratio-based approach effectively mitigates the impact of false-positive AMELY signals, therefore eliminating the need for time-consuming manual verification, and remains reliable even for samples with low peptide yields. Equipped with pre-constructed protein reference databases, *protSexInferer* provides a robust, standardized, and end-to-end solution for paleo-proteomic sex determination.

## Introduction

The accurate determination of the biological sex of ancient samples is a key facet of archaeological and anthropological research, and it is fundamental in the interpretation of sexual dimorphism features(Madupe et al., 2025), ancient demographic variations(García-Fernández et al., 2020; Goldberg et al., 2017; Gretzinger et al., 2022; Haak et al., 2015), and reconstructions of past social structures(Filiatreau, 2019; Fowler et al., 2022; J. Wang et al., 2025; Yüncü et al., 2025). And it is also valuable in forensic research(Dash et al., 2020; Krishan et al., 2016; Mikšík et al., 2023).

Morphological sex assessment relies on the analysis of osteological sexually dimorphic traits. This method, however, becomes less reliable when the address lacks critical morphological sex-diagnostic elements or features not fully developed(Krishan et al., 2016; MAYS, 2000; WALDRON, 1987). DNA-based molecular techniques determine sex by identifying sex chromosome-specific DNA markers or by quantifying the ratio of sequencing reads mapped to the X and Y chromosomes(Loreille et al., 2018; Mittnik et al., 2016; Skoglund et al., 2013). These approaches depend on the availability of adequately preserved DNA. The accurate sex-determination of remains with absent or undeveloped critical morphological sex-diagnostic features and unavailable DNA preservation is still an issue.

The development of mass spectrometry techniques, especially LC-MS/MS, to identify proteins in biological samples, enables the wide application of paleo-proteomics in sex identification(Gamble et al., 2024). This approach primarily relies on detecting sex-specific isoforms of the amelogenin gene — specifically, the AMELX, which is expressed on the X chromosome, and its Y-linked counterpart, AMELY(Fincham et al., 1991; Parker et al., 2019; Stewart et al., 2017). As the most abundant protein in dental enamel, amelogenin is particularly resistant to degradation and exogenous contamination owing to the highly mineralized nature of enamel(Castiblanco et al., 2015; Demarchi et al., 2016; Mazumder et al., 2014). Since amelogenin is more stable over time compared to proteins or DNA derived from less mineralized biological sources, the analysis of sex-specific amelogenin isoforms is feasible in poorly preserved ancient samples, even in some fragmented million-year-old fossils(Fong-Zazueta et al., 2025; Green et al., 2025; Taurozzi et al., 2024; T. Wang et al., 2025). Since it can be applied to subadult, fragmented, and endogenous DNA-free specimens, the paleo-proteomic amelogenin sex estimation has proven to be an effective and robust method for sex determination(Buonasera et al., 2020).

Despite the presence of over 20 amino acid differences between the human AMELY isoform and AMELX isoform 3(Parker et al., 2019), current workflows for paleo-proteomic sex estimation still rely on the detection of one (59M) or a few manually selected sexually dimorphic peptides(Stewart et al., 2017). Since the signal of AMELY-specific peptides is regarded as unambiguous evidence for a “female” individual in previous studies, it requires extensive manual verification by researchers specialized in paleo-proteomics to eliminate the risk of false positives AMELY signals in mass spectrometry database search(Adair et al., 2025; Cleland et al., 2024; Gowland et al., 2021; Lugli et al., 2019; Wasinger et al., 2019). Consequently, current paleo-proteomic sex estimation suffers from insufficient standardization and a lack of an integrated automatic pipeline, limiting its reproducibility, scalability, and overall robustness.

To address these limitations, we developed *protSexInferer*, an open-source Nextflow pipeline for paleo-proteomic sex estimation. Our pipeline determines biological sex by calculating the ratio of the number of AEMLY-specific peptides to all detected AMELX- and AMELY-specific amelogenin peptides (R_AMELY_). Sex classification thresholds are established based on the R_AMELY_ distribution in known sex samples and validated using independently published paleo-proteomic datasets. Furthermore, our pipeline integrates our pre-constructed amelogenin reference database and can parse the results of various search engines (PEAKS, pFind, MaxQuant, and DIA-NN). Its final results include information on all identified AMELX- and AMELY-specific peptides and sex assessment reports. Our results showed that *protSexInferer* provides a highly accurate, robust, end-to-end, and user-friendly solution for paleo-proteomic sex estimation.

## Results and discussions

### Workflow

#### Reference database construction

Before running *protSexInferer*, appropriate protein reference databases must be constructed or selected. The pipeline provides multiple pre-built databases optimized for different analytical contexts (Supplementary Text). By default, *protSexInferer* uses the “Hominidae enamel protein database”, including *Pongo*(Patramanis et al., 2023), *Gorilla*(Patramanis et al., 2023), *Pan*(Patramanis et al., 2023), *Homo*(Patramanis et al., 2023; Welker et al., 2020), *Gigantopithecus*(Welker et al., 2019), and *Paranthropus*(Madupe et al., 2025), designed for tooth enamel samples from unidentified primate specimens, with particular relevance to paleoanthropology research. The default enamel protein database comprises the following proteins: AHSG (FETUA), ALB (ALBU), AMBN, AMELX, AMELY, AMTN, COL17A1, ENAM, KLK4, MMP20, ODAM, and TUFT1. In addition, we provide alternative reference databases optimized for Neolithic archaeological samples, modern forensic specimens, and unidentified mammalian enamel samples (Supplementary Text). Users may select and apply the most appropriate reference database according to their specific research context. Using the same standardized reference databases ensures consistency in R_AMELY_ value calculations across studies.

#### Data input requirements

The *protSexInferer* accepts output files from commonly used protein search engines: PEAKS (v.11 or later; protein-peptides.csv) (Xin et al., 2022), MaxQuant (v.2.6.0.0 or later; evidence.txt) (Cox and Mann, 2008), pFind (v.3.2.1 or later; pFind.proteins) (Chi et al., 2018), and the DDA mode in DIA-NN (v.2.3.0 or later; report.parquet; currently available for *Homo* samples only) (Demichev et al., 2020). Database search must be performed by the user prior to running *protSexInferer*, with parameters optimized for ancient proteomic data as detailed in Supplementary Text.

#### Running *protSexInferer*

*protSexInferer* analysis proceeds through 3 main steps (Figure 1):

1. AMELX/AMELY-specific peptide classification
2. R_AMELY_ calculation and confidence interval estimation
3. Sex determination and report generation

**Figure 1:**
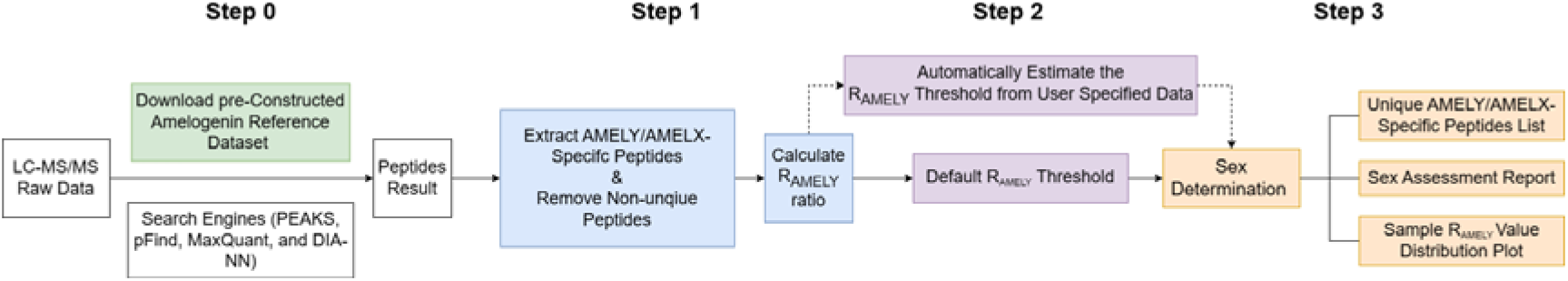
Overview of the *protSexInferer* Pipeline.

##### Step 1: AMELX/AMELY-specific peptide classification

The pipeline first parses database search results from the specified software platform. For non-PEAKS inputs, files are automatically converted to a PEAKS-compatible format. The pipeline removes modification information to obtain the base peptide sequence and merges identical peptide sequences to avoid duplicate counting. Then, the unique peptides matching the amelogenin proteins are extracted.

Classification of AMELY-specific and AMELX-specific peptides proceeds through two sequential filters. First, based on the database search results, peptides matching exclusively to AMELY (with no matches to AMELX) are considered putatively AMELY-specific, and vice versa for AMELX-specific peptides. Second, these putatively specific peptides are cross-validated against the complete reference database: any peptide showing identical matches to both AMELX and AMELY sequences—indicating origin from homologous regions—is excluded from quantification. Only peptides that pass both filters are considered authentic AMELX/Y-specific peptides for R_AMELY_ calculation.

##### Step 2: R_AMELY_ calculation and confidence interval estimation

For each sample, the pipeline calculates the R_AMELY_ ratio as: R_AMELY_ = n_AMELY_ /(n_AMELY_ + n_AMELX_), where n_AMELY_ represents the count of AMELY-specific peptides and n_AMELX_ represents the count of AMELX-specific peptides. Under the assumption that n_AMELY_ and n_AMELX_ follow a Bernoulli distribution (each detected peptide originates from either the X- or Y-chromosome), the pipeline computes 95% confidence intervals (CI) using the normal approximation: R_AMELY_ ± 1.960 × R_AMELY_ × (1-R_AMELY_) /(n_AMELY_ + n_AMELX_).

##### Step 3: Sex determination and report generation

Sex assignment is performed by comparing each sample’s R_AMELY_ value with established threshold ranges. By default, the pipeline uses pre-calculated thresholds specific to each search software, derived from known-sex reference individuals (Table S3). The female range is defined as R_AMELY_ values less than or equal to the upper limit of 95% CI of the observed maximum R_AMELY_ in reference females; the male range is defined as R_AMELY_ values greater than or equal to the lower limit of 95% CI of the observed minimum R_AMELY_ in reference males. Samples with R_AMELY_ values falling within one category’s range are assigned the corresponding sex; samples with values outside both ranges or overlapping the threshold boundary are reported as “Unknown Sex (U)”.

To facilitate downstream analysis of amelogenin peptides, the pipeline outputs comprehensive peptide information tables for each sample, listing all filtered amelogenin peptides. For each peptide, the table includes its sequence, classification status (“AMELX-unique”, “AMELY-unique”, or “Both”), and the summed “#Spec”, “Intensity”, and “Area” values. These results are accompanied by a R_AMELY_ value distribution plot with female and male threshold ranges shaded and samples colored by their assigned sex, and a final sex assignment report containing sample names, R_AMELY_ values with 95% confidence intervals (CI), and corresponding sex determinations.

#### Customization

While *protSexInferer* provides pre-built reference databases and optimized parameters for ancient proteomic data, users may customize these components when they are unsuitable for specific research contexts. Users can construct custom reference databases, provided that FASTA headers follow the required naming convention (Supplementary Text). When working with custom databases or search parameters, the default R_AMELY_ thresholds may not be appropriate. In such cases, users can re-estimate thresholds based on known-sex reference samples. The newly estimated thresholds can then be applied to unknown samples in subsequent analyses, offering flexibility for diverse sex determination experiments.

#### The effectiveness of *protSexInferer*’s sex identification

To establish the R_AMELY_ intervals for sex determination, we calculated the R_AMELY_ values for 76 reference samples of known sex(Lugli et al., 2019; Madupe et al., 2025; Parker et al., 2019; Rebay-Salisbury et al., 2020; Stewart et al., 2017). Detailed information on these reference individuals is listed in Table S1. We observed that, while the absolute R_AMELY_ ranges of the same LC-MS/MS raw data varied considerably after processing with different software, all R_AMELY_ values consistently formed two distinct clusters corresponding to sample sex (Figure 2). The R_AMELY_ thresholds for each search engine are provided in Table S3. Comparing across the 4 supported software platforms, we found MaxQuant yielded the lowest thresholds, suggesting more conservative detection of AMELY-specific signals with fewer false positives, while DIA-NN (the DDA mode) produced the highest thresholds, indicating more permissive peptide identification. Among all software, PEAKS exhibited the largest separation between the minimum R_AMELY_ of males and the maximum R_AMELY_ of females, offering the greatest discriminatory power. Based on these superior performances, we selected PEAKS as the default search engine and established its corresponding thresholds (male: R_AMELY_ > 0.088; female: R_AMELY_ < 0.055) for sex determination.

**Figure 2:**
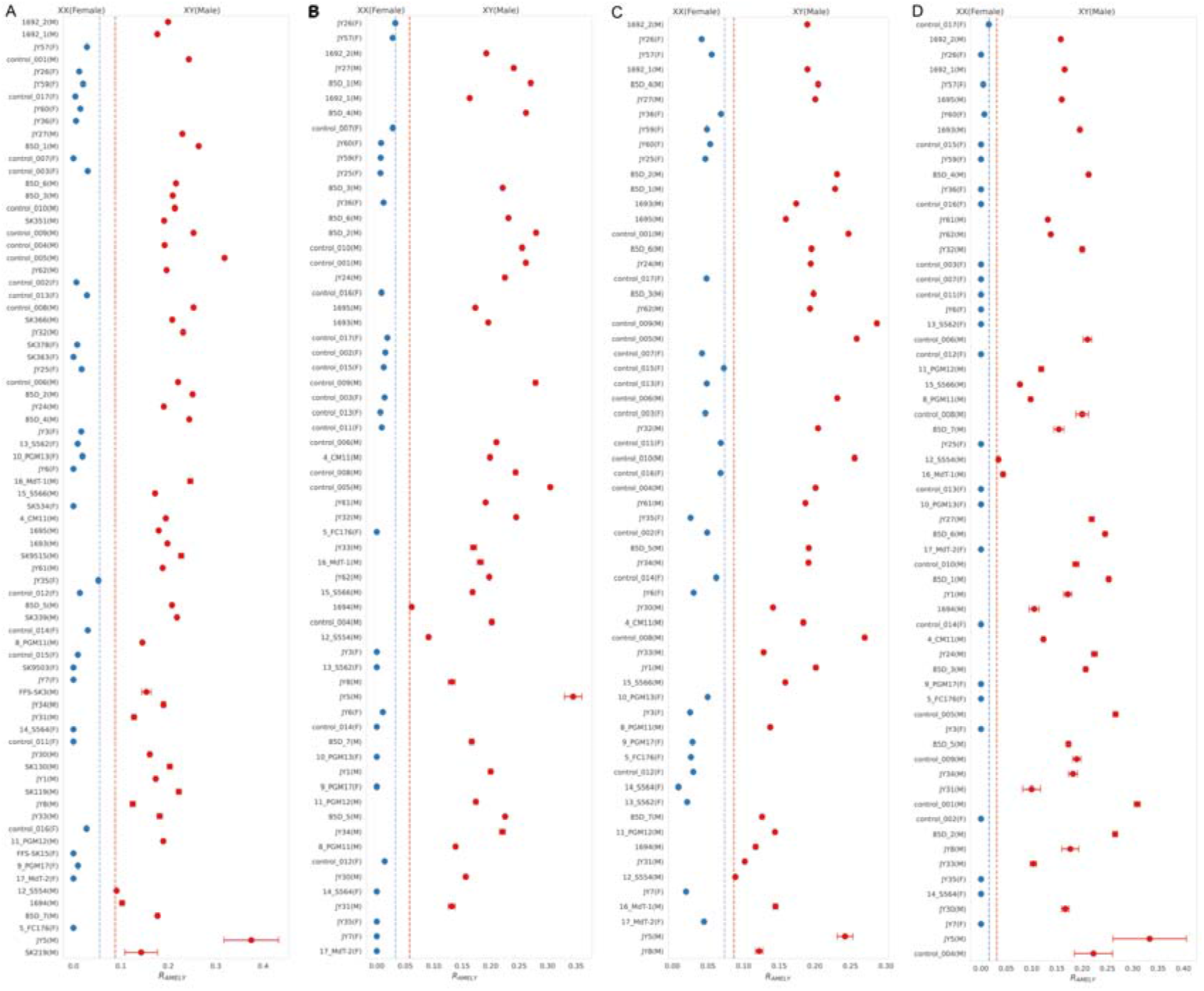
The estimated R_AMELY_ values for known sex reference samples. Values were calculated from the database search outputs of PEAKS (A), pFind (B), DIA-NN (C), and MaxQuant (D). We have indicated the prior determined sex of the published samples in parentheses after their sample names. The detailed information of these samples can be found in Table S1. We estimate the R_AMELY_ threshold to distinguish males and females based on the R_AMELY_ interval for each sex. The blue dashed line indicates the upper limit of 95% CI of the maximum R_AMELY_ for female individuals, and the red dashed line indicates the lower limit of 95% CI of the minimum R_AMELY_ for male individuals.

Subsequently, we applied the new pipeline to an independent validation dataset comprising 69 individuals whose sex had previously been determined via a proteomic-based method (Demeter et al., 2022; Gowland et al., 2021; Lugli et al., 2019; Madupe et al., 2025; Parker et al., 2019; Tsutaya et al., 2025; Welker et al., 2020), including both adult and non-adult individuals (also including the babies in gestation weeks), deciduous and permanent teeth, archaeological (up to ∼2Ma) and present-day specimens, sufficient and in sufficient enamel samples, and healthy and diseased teeth. Detailed information on these validation individuals is listed in Table S2. Our newly assigned sexes were consistent with prior determinations in almost all cases, regardless of the search engine used (Figure 3). This indicated that the R_AMELY_-based sex identification method is robust to the changes in database search software and specific parameters. Collectively, these results demonstrate that our novel pipeline serves as a robust and generalizable framework for paleo-proteomic sex determination. Notably, the individual JY63, previously morphologically assessed as male, was reclassified as female by our new method. This reclassification aligns with the results of the previous proteomic-based sex determination(Parker et al., 2019).

**Figure 3:**
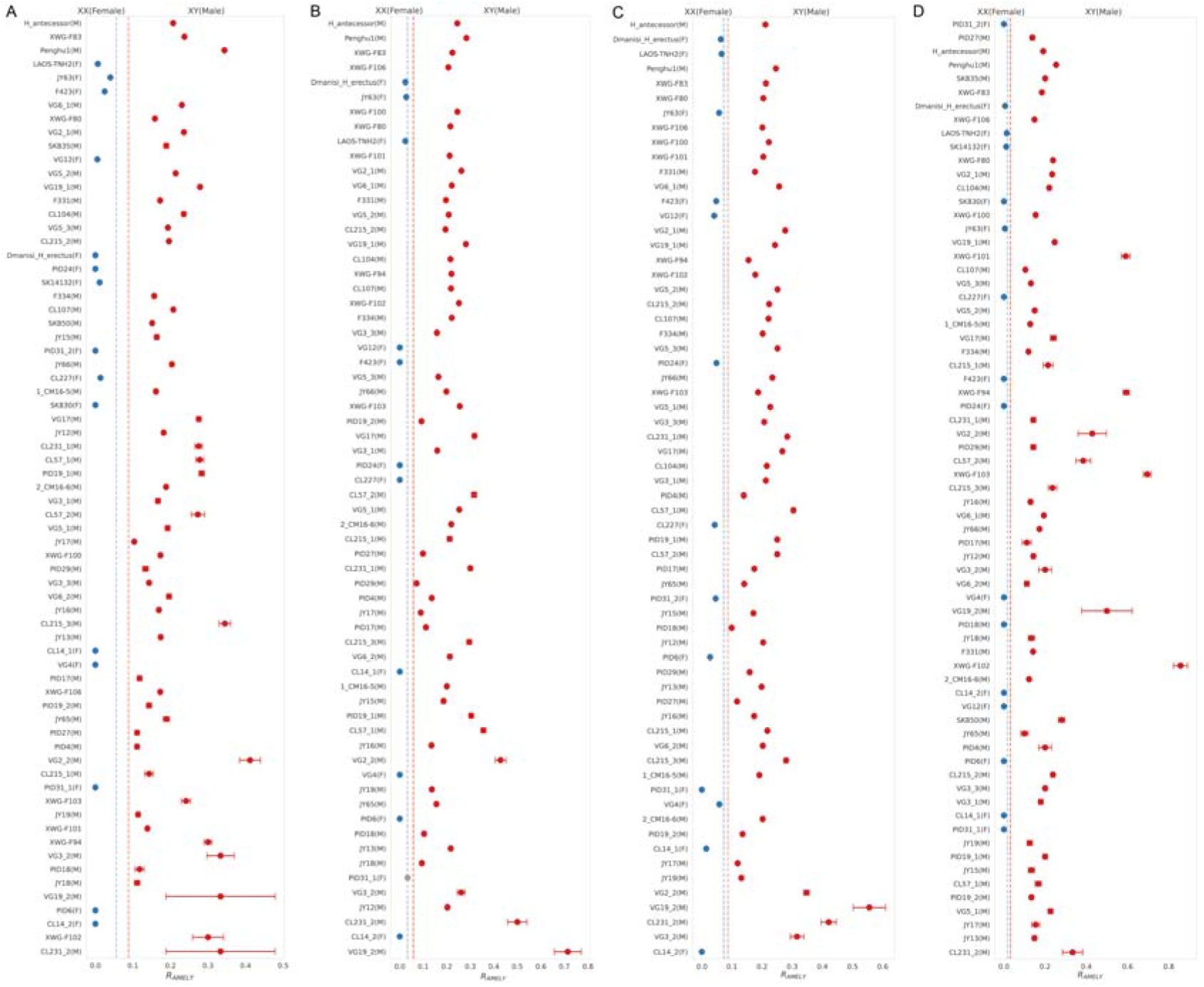
The estimated R_AMELY_ values for unknown sex validation samples and new reported samples from Xiawanggang Site. Values were calculated from the database search outputs of PEAKS (A), pFind (B), DIA-NN (C), and MaxQuant (D). The samples with R_AMELY_ values less than the female R_AMELY_ threshold are assigned as females (indicated by blue points), while samples with R_AMELY_ values larger than the male R_AMELY_ threshold are identified as males (indicated by red points).

#### The robustness of *protSexInferer* to the false positive AEMLY signals and low-data-amount samples

The signal of AMELY-specific peptides is commonly regarded as evidence of the presence of the Y chromosome, that is, a male individual. However, our results showed that although female individuals should theoretically not carry AMELY-specific peptides, AMELY-specific peptides were still detected in the database search results of some female individuals and persisted even after we repeatedly adjusted the reference datasets (Figure 4). And the false positive AMELY-specific peptide signals have been reported in several cases as well(Parker et al., 2019). These false positive signals also pose a risk of misclassifying females as males in sex determination approach that rely solely on the relative strength of the AMELY-59M and AMELX-60 peptides (Madupe et al., 2025).

**Figure 4:**
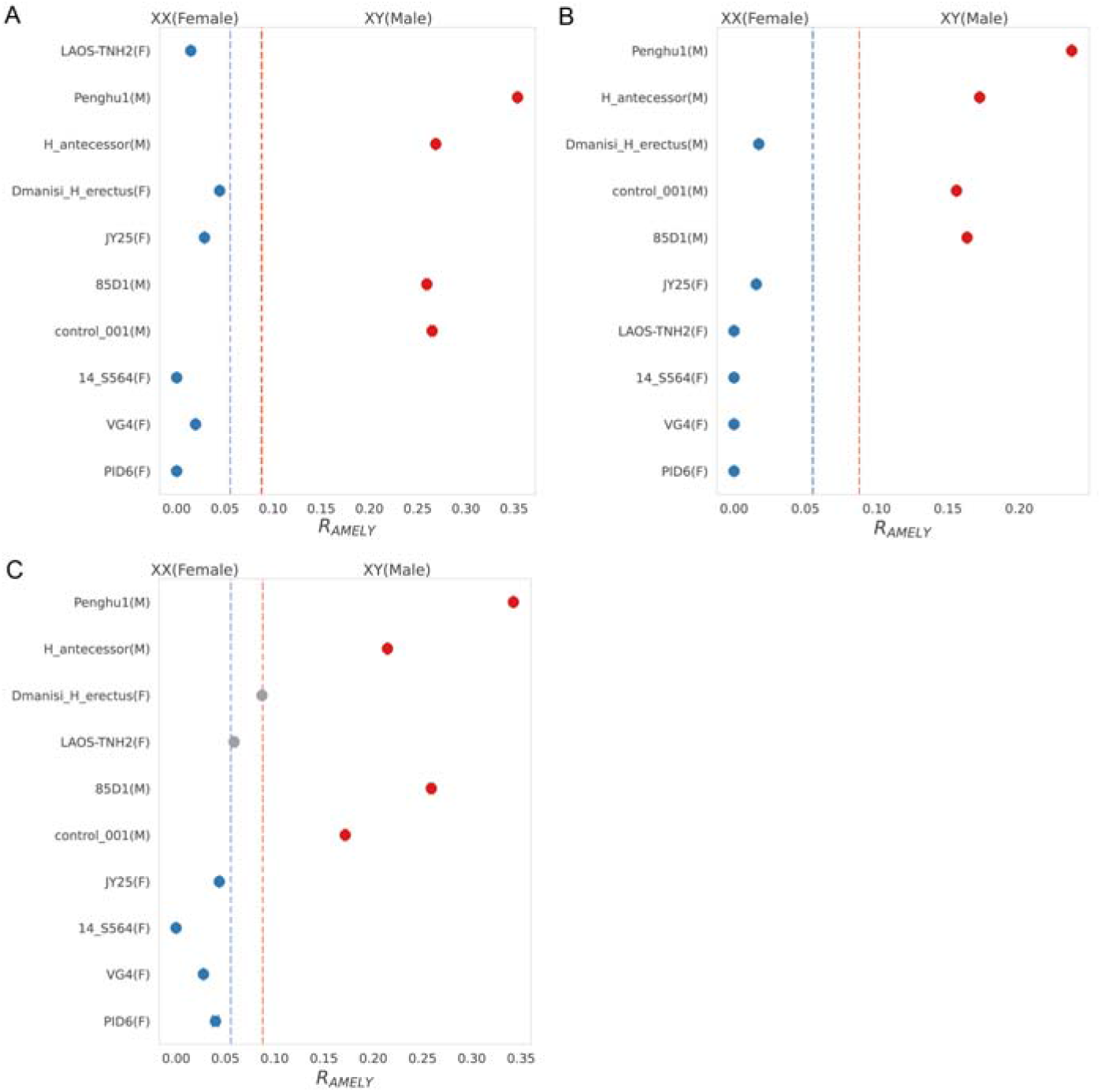
The estimated R_AMELY_ values for 10 randomly selected individuals calculated from PEAKS database search results using different reference databases. A) Searches using the “Human enamel proteins database” designed for human specimens. B) Searches using the “Hominidae AMEL database” designed for samples with high AMEL peptide content. C) Searches using the “Mammalian enamel proteins database” designed for unidentified mammalian samples. Dashed lines in the figure represent the R_AMELY_ thresholds calculated using the default “Hominidae enamel protein database”. Detailed information on each database can be found in Supplementary Text.

The false positive AMELY-specific peptide signals were in our expectation. In standard false discovery rate (FDR) reliability assessments, the “entrapment sequences” are introduced by adding random swapped/transversed mutations into the target human sequences, or by adding homologous proteins from genetically distant species (e.g., mouse or yeast). These “entrapment sequences” are expected to be detected at a ratio lower than the PSM FDR level. However, for the female specimens, as all AMELY sequences in the searching database have higher similarity than the standard “entrapment sequences” to the target AMELX sequences, the rate of false-positive AMELY identifications is expected to be modestly higher than the nominal PSM FDR in the sex determination analysis (0.010).

To evaluate the impact of these false positives, we applied the Madupe et al. method to results from 3 of the 4 supported search engines, as pFind was not supported for intensity information. For MaxQuant, which exhibited the lowest false positive AMELY rate, only one female individual was incorrectly classified as male. It was consistent with the lowest R_AMELY_ thresholds yielded by MaxQuant. However, for PEAKS and DIA-NN (DDA mode), the Madupe et al. method produced numerous false positive male assignments due to their more permissive peptide identification (Figure 5). In contrast, within our newly proposed R_AMELY_-based framework, these misclassified individuals can be reliably reassigned as female (Figure 3), eliminating the need for the time-consuming manual verification of all AMELY peptide identifications.

**Figure 5:**
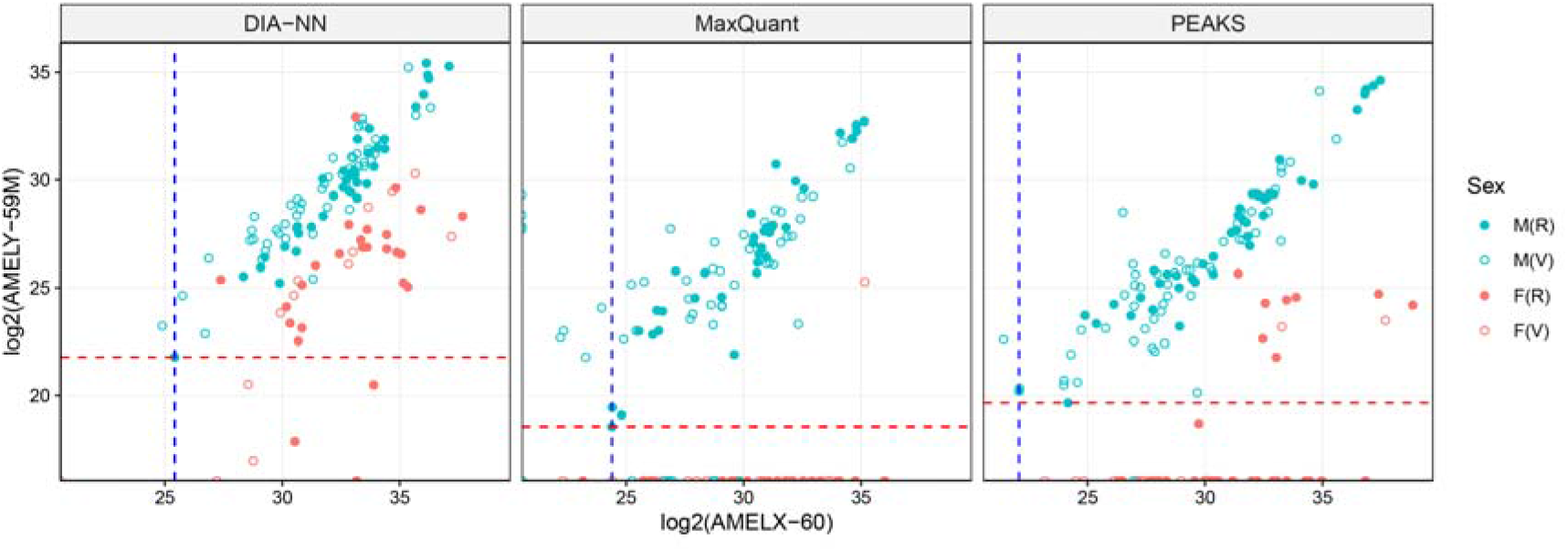
The relationship between the intensity of AMELY-59M and AMELX-60 peptides for reference and validation samples. M(R), M(V), F(R), and F(V) represent the results of reference male samples, validation male samples, reference female samples, and validation female samples, respectively. The blue vertical dashed line indicates the lowest log2(AMELX-60) value among the reference male individuals. The red horizontal dashed line indicates the lowest log2(AMELY-59) value among the reference male individuals. For the MaxQuant results, since the AMELY-59 signal was not detected in the JY33 individual, the second-lowest value (from the 16_MdT-1 individual) was used as the reference. The detailed information of all amelogenin peptides for these individuals was provided in the updated ProteomeXchange dataset (identifier: PXD072753).

Incorporating the full-length AMELY sequence substantially improved the sensitivity and accuracy of sex determination. In some instances where the intensities of 59M-related peptides were markedly low (less than 10% of typical levels), such as in samples VG3_2 and VG19_2(Gowland et al., 2021), or where the most typical AMELY-specific peptides (M(ox)IRPPY, *m*/*z* = 396.710; SMIRPPY, *m*/*z* = 432.230; SM(ox)IRPPY, *m*/*z* = 440.220) was partially or completely absent, additional AMELY-specific peptides enabled correct sex assignment. Those peptides would be helpful on badly preserved ancient specimens with no (or very few) 59M-related peptides detected (Figure 5), or on some basal mammals with different amino acids around AMELY-59. And the sex determination based on R_AMELY_ values could be effective and exact for both adult and non-adult individuals, and for both deciduous and permanent teeth.

To investigate how the total number of amelogenin peptides influences the estimated R_AMELY_ value in a sample, we evaluated their correlation. The results showed no strong correlation between the R_AMELY_ value and the total number of amelogenin peptides (R^2^ = 0.053, p = 0.004, for male individuals; R^2^ = 0.110, p = 0.007, for female individuals with R_AMELY_ value > 0; Figure S1). This indicated that, although the absolute numbers of amelogenin peptides fluctuated across ancient samples of varying ages and preservation conditions, the ratio of AEMLY-specific peptides was generally stable. As long as the signal from AMELY-specific peptides remains detectable, R_AMELY_ values can serve as a reliable reference for sex identification, even in enamel samples with insufficient mineralization or severe degradation. Furthermore, the estimated confidence intervals of R_AMELY_ values quantitatively reflected the uncertainty associated with the amount of data.

#### Application for sex determination of scattered teeth from Xiawanggang site (XWG)

The *protSexInferer* workflow was then applied for 8 randomly scattered teeth from the Xiawanggang site (Table S2). For all these specimens, the R_AMELY_ values calculated from the results of 4 search engines consistently exceeded the minimum male threshold (Figure 3), allowing them to be confidently classified as male.

#### Sex determination with other databases

To examine the effectiveness of the *protSexInferer* workflow using other databases, 3 different databases were tested on 10 randomly selected individuals. All 3 different databases showed large separation between the minimum R_AMELY_ of males and the maximum R_AMELY_ of females (Figure 4), and the results with databases smaller than the default “Homonidae enamel proteins database”, such as the “Human enamel proteins database” (Figure 4A) and the “Hominidae AMEL database” (Figure 4B), were similar to the default result, and could share the same thresholds. However, results with a larger “Mammalian enamel proteins database” did not fit the default thresholds, as we observed a notably higher false-positive rate (Figure 4C), possibly due to extra match errors introduced by the extra protein sequences, which are phylogenetically distant from the expected sample taxon. Thus, we strongly recommend using custom thresholds with any database larger than the default.

#### Limitations of our workflow

Our pipeline would be limited by the absence of the AMELY gene or expression in males(Mitchell et al., 2006; Turrina et al., 2011), though very rare. It would be limited by the random absence of AMELY-specific peptides in the enamel specimens, especially for the highly mineralized samples or insufficient sampling experiments. And this workflow is only applicable to mammalian species that possess both AMELX and AMELY isoforms, including primates, but excluding species such as mice, which have only a single isoform of AMEL(X) (Lau et al., 1989; Fincham et al., 1991).

## Methods and materials

### Pipeline framework and portability

*protSexInferer* is developed in the Nextflow programming language(Di Tommaso et al., 2017). Its built-in task scheduling and parallelization capabilities make the pipeline particularly suitable for large-scale sample batches. All software dependencies are managed through Nextflow’s Conda integration, which automatically handles environment configuration without requiring administrative privileges, ensuring adaptability to diverse computing platforms with minimal setup.

### Sample collection

We compiled a diverse enamel sample dataset for pipeline evaluation, comprising 76 samples with known sex and 69 samples with unknown sex from previously published studies (Table S1, Table S2). The 69 unknown-sex samples had prior sex determinations based on alternative proteomic methods. Additionally, we included 8 newly analyzed enamel samples from randomly scattered human teeth excavated from the Xiawanggang site, Henan, China (ca. 5,000 BP) (Henan Institute of Archaeology, 1989), for which sex information was previously undetermined.

### Protein extraction from the new Xiawanggang samples

An acid etching method was used to extract protein from tooth enamel(Stewart et al., 2017). Disposable toothbrushes were used to remove surface contaminants from a small area of enamel for etching. At the same time, the remaining teeth were wrapped with parafilm to prevent contact with any liquids. Before etching, the small enamel area was initially washed with 3% H_2_O_2_ for 30 seconds, followed by a rinse with ultrapure water. Approximately 100 µL of 5% (v/v) HCl was placed in the cap of a 1.5-mL microcentrifuge tube. A 2-minute etch was performed by immersing the etching region in the HCl solution, and the initial etch solution was discarded. A second etch, lasting 15 minutes, was carried out in the cap of another separate microcentrifuge tube, and the etch solution was retained. This second etch was repeated, and the etch solutions were combined and then desalted using C18 ZipTips (Thermo Fisher Scientific). After etching, the etched area was treated with 100 µL of 50 mM ammonium bicarbonate solution for 1 minute to neutralize the acid. It was then rinsed with ultrapure water for 30 seconds and dried. The protein peptides were eluted into a solution of 0.1% trifluoroacetic acid (TFA) and 80% acetonitrile (ACN). The peptide mixture was further divided into different aliquots and dried for Liquid chromatography-tandem mass spectrometry (LC-MS/MS) analysis. All sample preparation for the experiment was conducted in the dedicated clean room at the Molecular Paleontology Laboratory, IVPP of the Chinese Academy of Sciences in Beijing.

### Mass spectrometry data acquisition of the new Xiawanggang samples

The LC-MS/MS experiment was done with an Orbitrap Exploris 480 (Thermo) interfaced with an Easy n-LC 1200 HPLC system (Thermo) at Fudan University, Shanghai, China. The peptides were separated on a 75 μm id×25 cm analytical column, which was packed in-house using reversed-phase silica of 1.9 μm (Reprosil-Pur C18 AQ, Dr. Maisch GmbH). Buffer A was 0.1% formic acid in water, and Buffer B was 80% acetonitrile and 0.1% formic acid. An 80 min gradient was used with the following profile: 5-8% B, 2 min, at a flow rate of 200 nL/min; 8-44% B, 38 min, 200 nL/min; 44-70% B, 8 min, 200 nL/min; 70-100% B, 2min, 200 nL/min; 100% B, 10 min, 200 nL/min; 100-5% B, 2 min, 200 nL/min; 5% B, 2 min, 300 nL/min; 5-100% B, 6 min, 300 nl/min; 100% B, 10 min, 300 nL/min. Full MS scans were acquired for the first 65 min, after which the column was washed and re-equilibrated for 15 min without data acquisition. The full MS data acquisition was conducted across the range of *m/z* 350–1600, with a resolution of 60k at *m/z* 200. The AGC target was set to “Standard,” and the maximum injection time mode was set to “Auto”. The MS/MS spectra were acquired with a resolution of 15k at *m/z* 200, a maximum injection time of 30 ms, and a normalized collision energy of 30%. The AGC target was also set to “Standard”.

### Database search strategy

It is described in detail in Supplementary Text. The parameters were set to the default.

## Conclusion

In summary, our *protSexInferer* pipeline provides an automated approach for paleo-proteomic sex determination based on the R_AMELY_ ratio—the proportion of AMELY-specific peptides relative to total AMELX- and AMELY-specific peptides. By leveraging peptide ratios rather than binary signal detection, the method effectively eliminates false-positive AMELY identifications and obviates time-consuming manual verification. Unlike approaches targeting limited diagnostic sites (e.g., 59M), the pipeline incorporates all variant sites across AMELX and AMELY isoforms, fully utilizing available peptide data and maintaining high sensitivity even for low-quality samples. We showed that the R_AMELY_ values consistently discriminate sex across published reference and validation datasets. Compared to intensity-based methods, the ratio-based framework exhibits greater robustness against sporadic erroneous identifications. With integrated standardized databases and support for multiple search engine outputs, *protSexInferer* offers a robust, user-friendly, end-to-end solution for paleo-proteomic sex estimation.

## Acknowledgments

We thank our anonymous reviewers for their helpful comments. This study was supported by the Research Program of Chinese Academy of Sciences (XDA0460305), the National Science Foundation of China (32293192), the Chinese Academy of Sciences (CAS) (YSBR-019), and the Archaeological Talent Promotion Program of China (2024-278).

## Author contributions

Q.F. designed and supervised the research project. Z.W. processed the raw data. F.B. and Z.W. wrote the code, analyzed, and visualized the data. S.X. prepared the archaeological samples and materials. F.B., Z.W., and Q.F. did the data investigation, wrote the manuscript, discussed and revised the manuscript. Z.W. and F.B. wrote and prepared the supplementary materials. All authors discussed, critically revised, and approved the final version of the manuscript.

## Code and data availability

The source code of the *protSexInferer* pipeline is available on GitHub at https://github.com/QFuLab/protSexInferer with a manual. LC-MS/MS raw data, the estimated R_AMELY_ values, and filtered amelogenin peptides have been uploaded to the ProteomeXchange Consortium (http://proteomecentral.proteomexchange.org) with the data set identifier: PXD072753.

Reviewer access details: Log in to the PRIDE website using the following details: Project accession: PXD072753; Token: svedVIScJcs9; Alternatively, the reviewer can access the dataset by logging in to the PRIDE website using the following account details: Username; Password: oTDrhEs44J3Z.

## References

Adair, L.R., Lewis, M.E., Collins, M.J., Cramer, R., 2025. LAP-MALDI MS analysis of amelogenin from teeth for biological sex estimation. J Pharm Biomed Anal 255, 116599. 10.1016/j.jpba.2024.116599

Buonasera, T., Eerkens, J., de Flamingh, A., Engbring, L., Yip, J., Li, H., Haas, R., DiGiuseppe, D., Grant, D., Salemi, M., Nijmeh, C., Arellano, M., Leventhal, A., Phinney, B., Byrd, B.F., Malhi, R.S., Parker, G., 2020. A comparison of proteomic, genomic, and osteological methods of archaeological sex estimation. Sci Rep 10, 11897. 10.1038/s41598-020-68550-w

Castiblanco, G.A., Rutishauser, D., Ilag, L.L., Martignon, S., Castellanos, J.E., Mejía, W., 2015. Identification of proteins from human permanent erupted enamel. European Journal of Oral Sciences 123, 390–395. 10.1111/eos.12214

Chi, H., Liu, C., Yang, H., Zeng, W.-F., Wu, L., Zhou, W.-J., Wang, R.-M., Niu, X.-N., Ding, Y.-H., Zhang, Y., Wang, Z.-W., Chen, Z.-L., Sun, R.-X., Liu, T., Tan, G.-M., Dong, M.-Q., Xu, P., Zhang, P.-H., He, S.-M., 2018. Comprehensive identification of peptides in tandem mass spectra using an efficient open search engine. Nat Biotechnol 36, 1059–1061. 10.1038/nbt.4236

Cleland, T.P., McGuire, S.A., Beatrice, J.S., Moran, K.S., France, C.A.M., 2024. SPEED-E: A modified version of the sample preparation by Easy extraction and Digestion(-free) protocol for enamel-based sex estimation in archaeological remains. Journal of Archaeological Science 168, 106006. 10.1016/j.jas.2024.106006

Cox, J., Mann, M., 2008. MaxQuant enables high peptide identification rates, individualized p.p.b.-range mass accuracies and proteome-wide protein quantification. Nat Biotechnol 26, 1367–1372. 10.1038/nbt.1511

Dash, H.R., Rawat, N., Das, S., 2020. Alternatives to amelogenin markers for sex determination in humans and their forensic relevance. Mol Biol Rep 47, 2347–2360. 10.1007/s11033-020-05268-y

Demarchi, B., Hall, S., Roncal-Herrero, T., Freeman, C.L., Woolley, J., Crisp, M.K., Wilson, J., Fotakis, A., Fischer, R., Kessler, B.M., Rakownikow Jersie-Christensen, R., Olsen, J.V., Haile, J., Thomas, J., Marean, C.W., Parkington, J., Presslee, S., Lee-Thorp, J., Ditchfield, P., Hamilton, J.F., Ward, M.W., Wang, C.M., Shaw, M.D., Harrison, T., Domínguez-Rodrigo, M., MacPhee, R.D., Kwekason, A., Ecker, M., Kolska Horwitz, L., Chazan, M., Kröger, R., Thomas-Oates, J., Harding, J.H., Cappellini, E., Penkman, K., Collins, M.J., 2016. Protein sequences bound to mineral surfaces persist into deep time. eLife 5, e17092. 10.7554/eLife.17092

Demeter, F., Zanolli, C., Westaway, K.E., Joannes-Boyau, R., Duringer, P., Morley, M.W., Welker, F., Rüther, P.L., Skinner, M.M., McColl, H., Gaunitz, C., Vinner, L., Dunn, T.E., Olsen, J.V., Sikora, M., Ponche, J.-L., Suzzoni, E., Frangeul, S., Boesch, Q., Antoine, P.-O., Pan, L., Xing, S., Zhao, J.-X., Bailey, R.M., Boualaphane, S., Sichanthongtip, P., Sihanam, D., Patole-Edoumba, E., Aubaile, F., Crozier, F., Bourgon, N., Zachwieja, A., Luangkhoth, T., Souksavatdy, V., Sayavongkhamdy, T., Cappellini, E., Bacon, A.-M., Hublin, J.-J., Willerslev, E., Shackelford, L., 2022. A Middle Pleistocene Denisovan molar from the Annamite Chain of northern Laos. Nat Commun 13, 2557. 10.1038/s41467-022-29923-z

Demichev, V., Messner, C.B., Vernardis, S.I., Lilley, K.S., Ralser, M., 2020. DIA-NN: neural networks and interference correction enable deep proteome coverage in high throughput. Nat Methods 17, 41–44. 10.1038/s41592-019-0638-x

Di Tommaso, P., Chatzou, M., Floden, E.W., Barja, P.P., Palumbo, E., Notredame, C., 2017. Nextflow enables reproducible computational workflows. Nat Biotechnol 35, 316–319. 10.1038/nbt.3820

Filiatreau, C., 2019. Kinship-based social inequality in Bronze Age Europe. Science 12.

Fincham, A.G., Bessem, C.C., Lau, E.C., Pavlova, Z., Shuler, C., Slavkin, H.C., Snead, M.L., 1991. Human developing enamel proteins exhibit a sex-linked dimorphism. Calcif Tissue Int 48, 288–290. 10.1007/BF02556382

Fong-Zazueta, R., Krueger, J., Alba, D.M., Aymerich, X., Beck, R.M.D., Cappellini, E., Carrillo-Martin, G., Cirilli, O., Clark, N., Cornejo, O.E., Farh, K.K.-H., Ferrández-Peral, L., Juan, D., Kelley, J.L., Kuderna, L.F.K., Little, J., Orkin, J.D., Paterson, R.S., Pawar, H., Marques-Bonet, T., Lizano, E., 2025. Phylogenetic Signal in Primate Tooth Enamel Proteins and its Relevance for Paleoproteomics. Genome Biol Evol 17, evaf007. 10.1093/gbe/evaf007

Fowler, C., Olalde, I., Cummings, V., Armit, I., Büster, L., Cuthbert, S., Rohland, N., Cheronet, O., Pinhasi, R., Reich, D., 2022. A high-resolution picture of kinship practices in an Early Neolithic tomb. Nature 601, 584–587. 10.1038/s41586-021-04241-4

Gamble, J.A., Spicer, V., Hunter, M., Lao, Y., Hoppa, R.D., Pedersen, D.D., Wilkins, J.A., Zahedi, R.P., 2024. Advancing sex estimation from amelogenin: Applications to archaeological, deciduous, and fragmentary dental enamel. Journal of Archaeological Science: Reports 54, 104430. 10.1016/j.jasrep.2024.104430

García-Fernández, C., Font-Porterias, N., Kučinskas, V., Sukarova-Stefanovska, E., Pamjav, H., Makukh, H., Dobon, B., Bertranpetit, J., Netea, M.G., Calafell, F., Comas, D., 2020. Sex-biased patterns shaped the genetic history of Roma. Sci Rep 10, 14464. 10.1038/s41598-020-71066-y

Goldberg, A., Günther, T., Rosenberg, N.A., Jakobsson, M., 2017. Ancient X chromosomes reveal contrasting sex bias in Neolithic and Bronze Age Eurasian migrations. Proceedings of the National Academy of Sciences 114, 2657–2662. 10.1073/pnas.1616392114

Gowland, R., Stewart, N.A., Crowder, K.D., Hodson, C., Shaw, H., Gron, K.J., Montgomery, J., 2021. Sex estimation of teeth at different developmental stages using dimorphic enamel peptide analysis. Am J Phys Anthropol 174, 859–869. 10.1002/ajpa.24231

Green, D.R., Uno, K.T., Miller, E.R., Feibel, C.S., Aoron, E.E., Beck, C.C., Grossman, A., Kirera, F.M., Kirinya, M.M., Leakey, L.N., Liutkus-Pierce, C., Manthi, F.K., Ndiema, E.K., Nengo, I.O., Nyete, C., Rowan, J., Russo, G.A., Sanders, W.J., Smiley, T.M., Princehouse, P., Vitek, N.S., Cleland, T.P., 2025. Eighteen million years of diverse enamel proteomes from the East African Rift. Nature 643, 712–718. 10.1038/s41586-025-09040-9

Gretzinger, J., Sayer, D., Justeau, P., Altena, E., Pala, M., Dulias, K., Edwards, C.J., Jodoin, S., Lacher, L., Sabin, S., Vågene, Å.J., Haak, W., Ebenesersdóttir, S.S., Moore, K.H.S., Radzeviciute, R., Schmidt, K., Brace, S., Bager, M.A., Patterson, N., Papac, L., Broomandkhoshbacht, N., Callan, K., Harney, É., Iliev, L., Lawson, A.M., Michel, M., Stewardson, K., Zalzala, F., Rohland, N., Kappelhoff-Beckmann, S., Both, F., Winger, D., Neumann, D., Saalow, L., Krabath, S., Beckett, S., Van Twest, M., Faulkner, N., Read, C., Barton, T., Caruth, J., Hines, J., Krause-Kyora, B., Warnke, U., Schuenemann, V.J., Barnes, I., Dahlström, H., Clausen, J.J., Richardson, A., Popescu, E., Dodwell, N., Ladd, S., Phillips, T., Mortimer, R., Sayer, F., Swales, D., Stewart, A., Powlesland, D., Kenyon, R., Ladle, L., Peek, C., Grefen-Peters, S., Ponce, P., Daniels, R., Spall, C., Woolcock, J., Jones, A.M., Roberts, A.V., Symmons, R., Rawden, A.C., Cooper, A., Bos, K.I., Booth, T., Schroeder, H., Thomas, M.G., Helgason, A., Richards, M.B., Reich, D., Krause, J., Schiffels, S., 2022. The Anglo-Saxon migration and the formation of the early English gene pool. Nature 610, 112–119. 10.1038/s41586-022-05247-2

Haak, W., Lazaridis, I., Patterson, N., Rohland, N., Mallick, S., Llamas, B., Brandt, G., Nordenfelt, S., Harney, E., Stewardson, K., Fu, Q., Mittnik, A., Bánffy, E., Economou, C., Francken, M., Friederich, S., Pena, R.G., Hallgren, F., Khartanovich, V., Khokhlov, A., Kunst, M., Kuznetsov, P., Meller, H., Mochalov, O., Moiseyev, V., Nicklisch, N., Pichler, S.L., Risch, R., Rojo Guerra, M.A., Roth, C., Szécsényi-Nagy, A., Wahl, J., Meyer, M., Krause, J., Brown, D., Anthony, D., Cooper, A., Alt, K.W., Reich, D., 2015. Massive migration from the steppe was a source for Indo-European languages in Europe. Nature 522, 207–211. 10.1038/nature14317

Henan Institute of Archaeology, 1989. Xiawanggang in Xichuan (in Chinese). Cultural Relics Publishing House Press, Beijing.

Krishan, K., Chatterjee, P.M., Kanchan, T., Kaur, S., Baryah, N., Singh, R.K., 2016. A review of sex estimation techniques during examination of skeletal remains in forensic anthropology casework. Forensic Science International 261, 165.e1-165.e8. 10.1016/j.forsciint.2016.02.007

Lau, E.C., Mohandas, T.K., Shapiro, L.J., Slavkin, H.C., Snead, M.L., 1989. Human and mouse amelogenin gene loci are on the sex chromosomes. Genomics 4, 162–168. 10.1016/0888-7543(89)90295-4

Loreille, O., Ratnayake, S., Bazinet, A.L., Stockwell, T.B., Sommer, D.D., Rohland, N., Mallick, S., Johnson, P.L.F., Skoglund, P., Onorato, A.J., Bergman, N.H., Reich, D., Irwin, J.A., 2018. Biological Sexing of a 4000-Year-Old Egyptian Mummy Head to Assess the Potential of Nuclear DNA Recovery from the Most Damaged and Limited Forensic Specimens. Genes 9, 135. 10.3390/genes9030135

Lugli, F., Di Rocco, G., Vazzana, A., Genovese, F., Pinetti, D., Cilli, E., Carile, M.C., Silvestrini, S., Gabanini, G., Arrighi, S., Buti, L., Bortolini, E., Cipriani, A., Figus, C., Marciani, G., Oxilia, G., Romandini, M., Sorrentino, R., Sola, M., Benazzi, S., 2019. Enamel peptides reveal the sex of the Late Antique ‘Lovers of Modena.’ Sci Rep 9, 13130. 10.1038/s41598-019-49562-7

Madupe, P.P., Koenig, C., Patramanis, I., Rüther, P.L., Hlazo, N., Mackie, M., Tawane, M., Krueger, J., Taurozzi, A.J., Troché, G., Kibii, J., Pickering, R., Dickinson, M.R., Sahle, Y., Kgotleng, D., Musiba, C., Manthi, F., Bell, L., DuPlessis, M., Gilbert, C., Zipfel, B., Kuderna, L.F.K., Lizano, E., Welker, F., Kyriakidou, P., Cox, J., Mollereau, C., Tokarski, C., Blackburn, J., Ramos-Madrigal, J., Marques-Bonet, T., Penkman, K., Zanolli, C., Schroeder, L., Racimo, F., Olsen, J.V., Ackermann, R.R., Cappellini, E., 2025. Enamel proteins reveal biological sex and genetic variability in southern African Paranthropus. Science 388, 969–973. 10.1126/science.adt9539

Mays, S., 2000. Sex determination in skeletal remains. Human Osteology in Archeology and Forensic Science 117–130.

Mazumder, P., Prajapati, S., Lokappa, S.B., Gallon, V., Moradian-Oldak, J., 2014. Analysis of co-assembly and co-localization of ameloblastin and amelogenin. Front. Physiol. 5. 10.3389/fphys.2014.00274

Mikšík, I., Morvan, M., Brůžek, J., 2023. Peptide analysis of tooth enamel – A sex estimation tool for archaeological, anthropological, or forensic research. Journal of Separation Science 46, 2300183. 10.1002/jssc.202300183

Mitchell, R.J., Kreskas, M., Baxter, E., Buffalino, L., Van Oorschot, R.A.H., 2006. An investigation of sequence deletions of amelogenin (AMELY), a Y-chromosome locus commonly used for gender determination. Annals of Human Biology 33, 227–240. 10.1080/03014460600594620

Mittnik, A., Wang, C.-C., Svoboda, J., Krause, J., 2016. A Molecular Approach to the Sexing of the Triple Burial at the Upper Paleolithic Site of Dolní Věstonice. PLOS ONE 11, e0163019. 10.1371/journal.pone.0163019

Parker, G.J., Yip, J.M., Eerkens, J.W., Salemi, M., Durbin-Johnson, B., Kiesow, C., Haas, R., Buikstra, J.E., Klaus, H., Regan, L.A., Rocke, D.M., Phinney, B.S., 2019. Sex estimation using sexually dimorphic amelogenin protein fragments in human enamel. Journal of Archaeological Science 101, 169–180. 10.1016/j.jas.2018.08.011

Patramanis, I., Ramos-Madrigal, J., Cappellini, E., Racimo, F., 2023. PaleoProPhyler: a reproducible pipeline for phylogenetic inference using ancient proteins. Peer Community Journal 3. 10.24072/pcjournal.344

Rebay-Salisbury, K., Janker, L., Pany-Kucera, D., Schuster, D., Spannagl-Steiner, M., Waltenberger, L., Salisbury, R.B., Kanz, F., 2020. Child murder in the Early Bronze Age: proteomic sex identification of a cold case from Schleinbach, Austria. Archaeol Anthropol Sci 12, 265. 10.1007/s12520-020-01199-8

Skoglund, P., Storå, J., Götherström, A., Jakobsson, M., 2013. Accurate sex identification of ancient human remains using DNA shotgun sequencing. Journal of Archaeological Science 40, 4477–4482. 10.1016/j.jas.2013.07.004

Stewart, N.A., Gerlach, R.F., Gowland, R.L., Gron, K.J., Montgomery, J., 2017. Sex determination of human remains from peptides in tooth enamel. Proc Natl Acad Sci U S A 114, 13649–13654. 10.1073/pnas.1714926115

Taurozzi, A.J., Rüther, P.L., Patramanis, I., Koenig, C., Sinclair Paterson, R., Madupe, P.P., Harking, F.S., Welker, F., Mackie, M., Ramos-Madrigal, J., Olsen, J.V., Cappellini, E., 2024. Deep-time phylogenetic inference by paleoproteomic analysis of dental enamel. Nat Protoc 19, 2085–2116. 10.1038/s41596-024-00975-3

Tsutaya, T., Sawafuji, R., Taurozzi, A.J., Fagernäs, Z., Patramanis, I., Troché, G., Mackie, M., Gakuhari, T., Oota, H., Tsai, C.-H., Olsen, J.V., Kaifu, Y., Chang, C.-H., Cappellini, E., Welker, F., 2025. A male Denisovan mandible from Pleistocene Taiwan. Science 388, 176–180. 10.1126/science.ads3888

Turrina, S., Filippini, G., Voglino, G., De Leo, D., 2011. Two additional reports of deletion on the short arm of the Y chromosome. Forensic Science International: Genetics 5, 242–246. 10.1016/j.fsigen.2010.10.015

Waldron, T., 1987. The Relative Survival of the Human Skeleton□: Implications for Palaeopathology. Death, Decay and Reconstruction□: Approaches to Archaeology and Forensic Science 55–64.

Wang, J., Yan, S., Li, Z., Zan, J., Zhao, Y., Zhao, J., Chen, K., Wang, X., Ji, T., Zhang, C., Yang, T., Zhang, T., Qiao, R., Guo, M., Rao, Z., Zhang, J., Wang, G., Ran, Z., Duan, C., Zhang, F., Song, Y., Wu, X., Mace, R., Sun, B., Pang, Y., Huang, Y., Zhang, H., Ning, C., 2025. Ancient DNA reveals a two-clanned matrilineal community in Neolithic China. Nature 643, 1304–1311. 10.1038/s41586-025-09103-x

Wang, T., Yang, M.A., Zhu, Z., Ma, M., Shi, H., Speidel, L., Min, R., Yuan, H., Jiang, Z., Hu, C., Li, X., Zhao, D., Bai, F., Cao, P., Liu, F., Dai, Q., Feng, X., Yang, R., Wu, X., Liu, X., Zhang, M., Ping, W., Liu, Y., Wan, Y., Yang, F., Zhou, R., Kang, L., Dong, G., Stoneking, M., Fu, Q., 2025. Prehistoric genomes from Yunnan reveal ancestry related to Tibetans and Austroasiatic speakers. Science 388, eadq9792. 10.1126/science.adq9792

Wasinger, V.C., Curnoe, D., Bustamante, S., Mendoza, R., Shoocongdej, R., Adler, L., Baker, A., Chintakanon, K., Boel, C., Tacon, P.S.C., 2019. Analysis of the Preserved Amino Acid Bias in Peptide Profiles of Iron Age Teeth from a Tropical Environment Enable Sexing of Individuals Using Amelogenin MRM. PROTEOMICS 19, 1800341. 10.1002/pmic.201800341

Welker, F., Ramos-Madrigal, J., Gutenbrunner, P., Mackie, M., Tiwary, S., Rakownikow Jersie-Christensen, R., Chiva, C., Dickinson, M.R., Kuhlwilm, M., de Manuel, M., Gelabert, P., Martinón-Torres, M., Margvelashvili, A., Arsuaga, J.L., Carbonell, E., Marques-Bonet, T., Penkman, K., Sabidó, E., Cox, J., Olsen, J.V., Lordkipanidze, D., Racimo, F., Lalueza-Fox, C., Bermúdez de Castro, J.M., Willerslev, E., Cappellini, E., 2020. The dental proteome of Homo antecessor. Nature 580, 235–238. 10.1038/s41586-020-2153-8

Welker, F., Ramos-Madrigal, J., Kuhlwilm, M., Liao, W., Gutenbrunner, P., de Manuel, M., Samodova, D., Mackie, M., Allentoft, M.E., Bacon, A.-M., Collins, M.J., Cox, J., Lalueza-Fox, C., Olsen, J.V., Demeter, F., Wang, W., Marques-Bonet, T., Cappellini, E., 2019. Enamel proteome shows that Gigantopithecus was an early diverging pongine. Nature 576, 262–265. 10.1038/s41586-019-1728-8

Xin, L., Qiao, R., Chen, X., Tran, H., Pan, S., Rabinoviz, S., Bian, H., He, X., Morse, B., Shan, B., Li, M., 2022. A streamlined platform for analyzing tera-scale DDA and DIA mass spectrometry data enables highly sensitive immunopeptidomics. Nat Commun 13, 3108. 10.1038/s41467-022-30867-7

Yüncü, E., Doğu, A.K., Kaptan, D., Kılıç, M.S., Mazzucato, C., Güler, M.N., Eker, E., Katırcıoğlu, B., Chyleński, M., Vural, K.B., Sağlıcan, E., Atağ, G., Bozkurt, D., Pearson, J., Sevkar, A., Altınışık, N.E., Milella, M., Karamurat, C., Aktürk, Ş., Yurttaş, E.D., Yıldız, N., Koptekin, D., Yorulmaz, S., Kazancı, D.D., Aydoğan, A., Gürün, K., Schotsmans, E.M.J., Anvari, J., Rosenstock, E., Byrnes, J., Biehl, P.F., Orton, D., Lagerholm, V.K., Gemici, H.C., Vasic, M., Marciniak, A., Atakuman, Ç., Erdal, Y.S., Kırdök, E., Pilloud, M., Larsen, C.S., Haddow, S.D., Götherström, A., Knüsel, C.J., Özer, F., Hodder, I., Somel, M., 2025. Female lineages and changing kinship patterns in Neolithic Çatalhöyük. Science 388, eadr2915. 10.1126/science.adr2915

