## Supplementary Text for "Rapid and robust sex determination from ancient enamel proteomes using *protSexInferer*"

**Overview**

protSexInferer estimates the sex of ancient samples based on their R_AMELY_ value, which is the ratio of the number of ancient samples’ AMELY-specific peptides to the total number of AMELY- and AMELX-specific peptides.

This pipeline uses the protein database searching results as the input data, and it supports popular protein search software, such as PEAKS^1^, MaxQuant^2^, pFind^3^, and the DDA mode of DIA-NN^4^. (Note: DIA-NN is only available for *Homo* samples now. We would continually add supporting software in the future, like MSFragger^5^, in our plan.) The default R_AMELY_ threshold for female and male of each software was provided within the pipeline, and the user can also re-estimate the R_AMELY_ threshold of their specific samples using this pipeline.

The pipeline will output the information of all filtered AMELY- and AMELX-specific peptides, the input samples’ R_AMELY_ value and confidence interval, the final sex estimation report, and the R_AMELY_ distribution plot.

**How to prepare input data**

1) Prepare the protein reference database

The default “Hominidae enamel protein database” (hepdb.fasta) contains AHSG, ALB, AMBN, AMELX, AMELY, AMTN, COL17A1, ENAM, MMP20, ODAM, KLK4, and TUFT1 of all Hominidae species from the PaleoProPhyler database^6^ (lack of KLK4 and TUFT1; AHSG was wrong and thus not included), UniProt, our translation from published genomes ^7-13^, and mutated sequences from ancient proteomes ^14-17^.

There were other filtered reference databases to choose: a) a “Human enamel proteins database”, optimized for tooth enamel samples from human specimens, especially for Neolithic archaeology research or forensic research (hhepdb.fasta); b) a “Hominidae AMEL database”, for samples with high AMEL level, or only AMEL (hxy.fasta); c) a “Mammalian enamel proteins database”, designed for tooth enamel samples from unidentified mammalian specimens (mepdb.fasta).

If the provided pre-built database does not meet the requirements, users can also build their own reference database. However, because the pipeline assigns sequences to genes based on their FASTA header annotations, sequence names in the reference database must follow a defined naming convention. Specifically, the gene identity must be indicated either by including a gene tag in the FASTA description using the format GN=XX (e.g., >Seq1 [GN=AMELY]), or by placing the gene name at the beginning of the sequence identifier, separated from the rest by a hyphen (-) or underscore (_) (e.g., >AMELY_Pabe1v2 or >AMELY-202_Pabe1v2).

**Note:** If the user uses a non-default pre-built reference database (a, b, c above) or a user-built reference database, the R_AMELY_ threshold used for sex identification needs to be re-estimated.

2) Searched the LC-MS/MS raw data against the reference database

We recommend using these parameters for Orbitrap HCD raw data in DDA mode:

a) PEAKS (v.11 or later): Precursor Mass Error Tolerance = 10 ppm, Fragment Mass Error Tolerance = 0.02 Da, “None” Enzyme, Peptide Length = 6-45, open “Deep Learning Boost”, no Fixed Modification, including Deamidation (NQ), Hydroxylation(P), Oxidation (M), Phosphorylation (STY), Pyro-glu from E, Pyro-glu from Q as Variable Modifications, and Max Variable PTM Per Peptide = 3, “Use Database Search Parameters” for *de novo*, open “PEAKS PTM”, open “SPIDER”, filter PSM with 1% FDR, filter protein with -10LgP ≥ 20, and filter *de novo* Only ALC ≥ 50%;

**Note:** For CID raw data with a higher mass error level, please use: Precursor Mass Error Tolerance = 20 ppm, Fragment Mass Error Tolerance = 0.5 Da; or any other suggested value of the user’s LC-MS/MS instrument.

b) MaxQuant (v.2.6.0.0 or later): Unspecific Digestion, no Fixed Modification, including Deamidation (NQ), Oxidation(P), Oxidation (M), Phosphorylation (STY), and Max Variable PTM Per Peptide = 3, filter PSM with 1% FDR, filter protein with 10% FDR, open “second peptides”, other parameters were as default;

c) pFind (v.3.2.1 or later): msmstype=PF1/2, MS1 mass tolerance = 10 ppm, MS2 mass tolerance = 0.02 Da, NoEnzyme U_C, Peptide Length = 6-45, mass range = 360-4500 Da, no Fixed Modification, including Deamidated[N], Deamidated[Q], Gln->pyro-Glu[AnyN-termQ], Glu->pyro-Glu[AnyN-termE], Oxidation[M], Phospho[S], Phospho[T], Phospho[Y], and Oxidation[P] as Variable Modifications (for ancient samples, Oxidation[W], Dioxidation[M], Arg->Orn[R], Dioxidation[W], Trp->Kynurenin[W] were also included as Variable Modifications), and Max Variable PTM Per Peptide = 3, open “Open search”, filter PSM with 1% FDR, filter protein with 10% FDR;

d) DIA-NN (v.2.3.0 or later): --dda --verbose 1 --cut ** --missed-cleavages 100 --qvalue 0.01 --var-mod UniMod:7,0.984016,NQ --var-mod UniMod:27,-18.010565,E --var-mod UniMod:28,-17.026549,Q --var-mod UniMod:35,15.994915,MPW (W only for ancient samples) --var-mod UniMod:425,31.989829,MW (only for ancient samples) --var-mod UniMod:21,79.966331,STY --matrices --ignore-decoys --high-acc --smart-profiling --xic --xic-theoretical-fr --duplicate-proteins --export-quant --no-swissprot --pg-level 0 --predictor.

3) Collected the database searching results

We need different files from the results of different database searching software: a) PEAKS: protein-peptides.csv; b) MaxQuant: evidence.txt; c) pFind: pFind.proteins; d) DIA-NN: report.parquet.

**Parameters**

--inputFile: The path of the input CSV table. Its first column is the sample name, and its second column is the path of the corresponding database searching result file. If the --findrAMELYThreshold’ parameter is set, the input CSV table needs a third column, which is the sex of the input samples.

--databaseSearchingSoftware: The name of database searching software. It can only be one of these values: PEAKS, MaxQuant, DIA-NN, or pFind. The default value is PEAKS.

--findrAMELYThreshold: If this parameter is set, the pipeline will estimate the R_AMELY_ threshold based on the input known sex reference samples.

--femaleMaxrAMELY: User can manually set the maximum R_AMELY_ of female using this parameter. Mainly works with the ‘--findrAMELYThreshold’ parameter.

--maleMinrAMELY: User can manually set the minimum R_AMELY_ of male using this parameter. Mainly works with the ‘--findrAMELYThreshold’ parameter.

--autoBLAST: If this parameter is set, all filtered AMELY-specific peptides longer than the length specified by the ‘--minimumBLASTLength’ parameter will go through automatic BLAST, and any peptides which 100% matches to the non-amelogenin gene will be removed. This step is time consuming, and it is turn off in default.

--withoutBLASTdb: If this parameter is set, the pipeline will download the whole BLAST reference databases into the ‘BLAST_Database’ directory. This step is time-consuming and involves heavy network usage, and it is turned off by default.

--minimumBLASTLength: The minimum length of peptide that went through BLAST. Only works with the ‘--autoBLAST’ parameter. The default value is 8. Not recommend using a value lower than 6, because the short segment has a high possibility to match the non-amelogenin gene, even if it’s an authentic AMELY-specific peptide.

--outDir: The path of the directory saving all output files.

**Dependencies**

This pipeline is based on Nextflow. It’s dependent on publicly available software or packages that were listed in the ‘nextflow.config’ file. And all of the software and packages can be installed by Nextflow’s built-in support for Conda environments.

**Usage**

1) Using the default R_AMELY_ threshold

Run the analysis using the command `nextflow run protSexInferer.nf --inputFile input.csv --outDir output `.

The information table of filtered AMELY- and AMELX-specific peptides will be saved under the ‘output/classified_peptides’ directory. The final sex estimation report will be saved as the ‘output/Sex_assessment_report.csv’ file. The distribution of samples’ R_AMELY_ value will be saved as the ‘output/rAMELY_protSexInferer.pdf’ file.

2) Using the re-estimated R_AMELY_ threshold

Firstly, run the command ` nextflow run protSexInferer.nf --inputFile input.csv --outDir newThreshold --findrAMELYThreshold` to estimate the R_AMELY_ value based on user-provided known sex reference samples. The R_AMELY_ threshold will be reported in the ‘newThreshold/rAMELY_threshold.txt’ file.

Then, estimate the sex of input samples using the new estimated R_AMELY_ threshold assigned by the ‘--femaleMaxrAMELY’ and ‘--maleMinrAMELY’ parameters. The example command is `nextflow run protSexInferer.nf --inputFile input.csv --outDir output --femaleMaxrAMELY 0.03 --maleMinrAMELY 0.08`.

**REFERENCE**

1	Xin, L. *et al.* A streamlined platform for analyzing tera-scale DDA and DIA mass spectrometry data enables highly sensitive immunopeptidomics. *Nat Commun* **13**, 3108, doi:10.1038/s41467-022-30867-7 (2022).

2	Cox, J. & Mann, M. MaxQuant enables high peptide identification rates, individualized p.p.b.-range mass accuracies and proteome-wide protein quantification. *Nat Biotechnol* **26**, 1367-1372, doi:10.1038/nbt.1511 (2008).

3	Chi, H. *et al.* Comprehensive identification of peptides in tandem mass spectra using an efficient open search engine. *Nat Biotechnol*, doi:10.1038/nbt.4236 (2018).

4	Demichev, V., Messner, C. B., Vernardis, S. I., Lilley, K. S. & Ralser, M. DIA-NN: neural networks and interference correction enable deep proteome coverage in high throughput. *Nat Methods* **17**, 41-44, doi:10.1038/s41592-019-0638-x (2020).

5	Kong, A. T., Leprevost, F. V., Avtonomov, D. M., Mellacheruvu, D. & Nesvizhskii, A. I. MSFragger: ultrafast and comprehensive peptide identification in mass spectrometry-based proteomics. *Nat Methods* **14**, 513-520, doi:10.1038/nmeth.4256 (2017).

6	Patramanis, I., Ramos-Madrigal, J., Cappellini, E. & Racimo, F. PaleoProPhyler: a reproducible pipeline for phylogenetic inference using ancient proteins. *Peer Community Journal* **3**, doi:10.24072/pcjournal.344 (2023).

7	Mafessoni, F. *et al.* A high-coverage Neandertal genome from Chagyrskaya Cave. *Proc Natl Acad Sci U S A* **117**, 15132-15136, doi:10.1073/pnas.2004944117 (2020).

8	Kronenberg, Z. N. *et al.* High-resolution comparative analysis of great ape genomes. *Science* **360**, eaar6343, doi:doi:10.1126/science.aar6343 (2018).

9	Bergström, A. *et al.* Insights into human genetic variation and population history from 929 diverse genomes. *Science* **367**, eaay5012, doi:doi:10.1126/science.aay5012 (2020).

10	Prüfer, K. *et al.* A high-coverage Neandertal genome from Vindija Cave in Croatia. *Science* **358**, 655-658, doi:doi:10.1126/science.aao1887 (2017).

11	Prufer, K. *et al.* The complete genome sequence of a Neanderthal from the Altai Mountains. *Nature* **505**, 43-49, doi:10.1038/nature12886 (2014).

12	Meyer, M. *et al.* A High-Coverage Genome Sequence from an Archaic Denisovan Individual. *Science* **338**, 222-226, doi:doi:10.1126/science.1224344 (2012).

13	Lalueza-Fox, C., Rosas, A. & de la Rasilla, M. Palaeogenetic research at the El Sidron Neanderthal site. *Ann Anat* **194**, 133-137, doi:10.1016/j.aanat.2011.01.014 (2012).

14	Tsutaya, T. *et al.* A male Denisovan mandible from Pleistocene Taiwan. *Science* **388**, 176-180, doi:10.1126/science.ads3888 (2025).

15	Madupe, P. P. *et al.* Enamel proteins reveal biological sex and genetic variability in southern African *Paranthropus*. *Science* **388**, 969-973, doi:10.1126/science.adt9539 (2025).

16	Welker, F. *et al.* The dental proteome of *Homo antecessor*. *Nature* **580**, 235-238, doi:10.1038/s41586-020-2153-8 (2020).

17	Welker, F. *et al.* Enamel proteome shows that *Gigantopithecus* was an early diverging pongine. *Nature* **576**, 262-265, doi:10.1038/s41586-019-1728-8 (2019).

Supplementary Figures

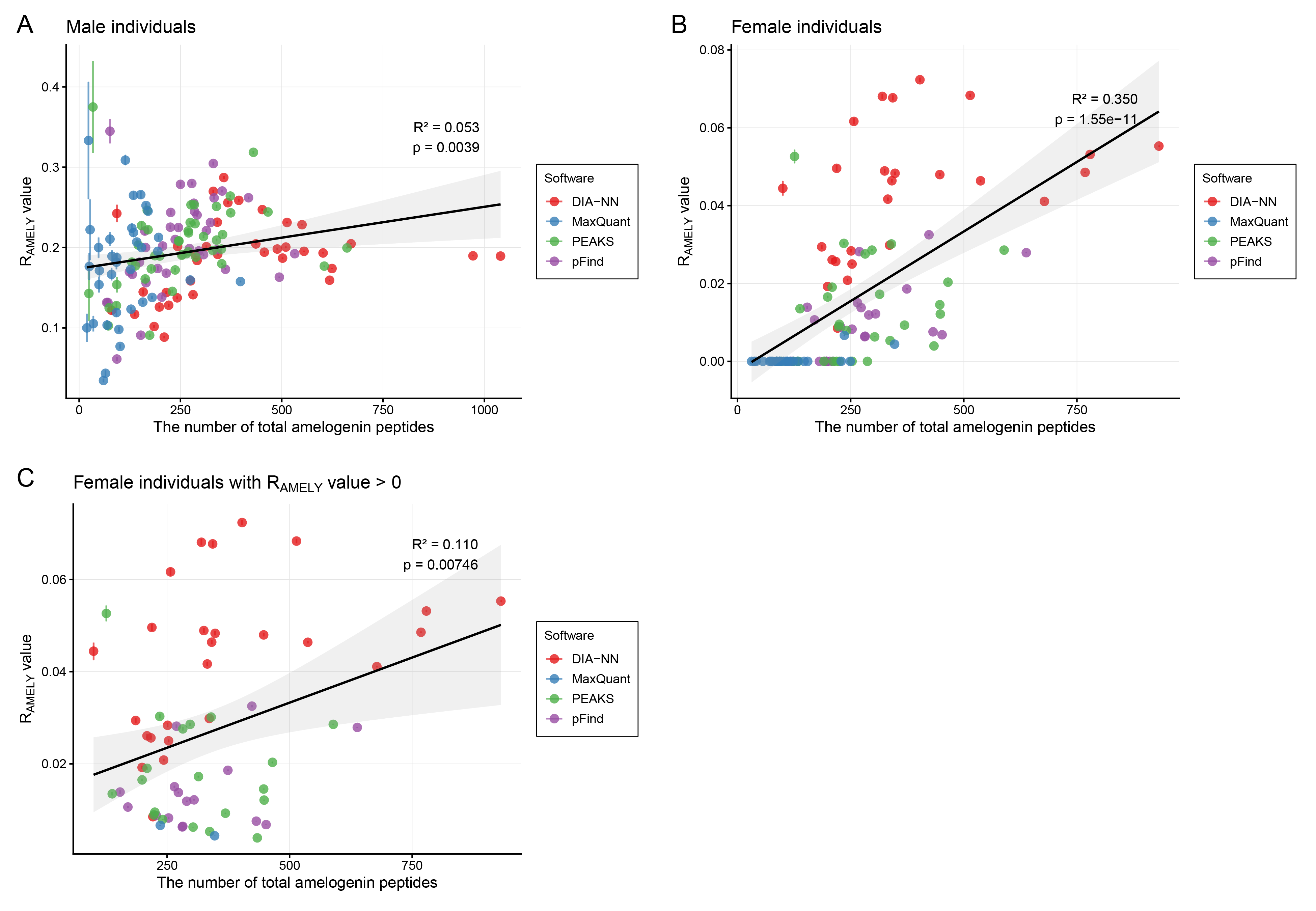

**Figure S1: The correlation between the number of total amelogenin peptides (x-axis) and R_AMELY_ values (y-axis) for all male samples (A), all female samples (B), and female samples with R_AMELY_ value > 0 (C).** Different colors represent the results from different database searching software. The R² and p values are annotated in the figure.
